# Bleomycin induces active-centromere damage and cytoplasmic mislocalization of centromeric chromatin in fibroblasts

**DOI:** 10.1101/2025.08.22.671795

**Authors:** Azait Imtiaz, Mohammad Waseem, Hudson O’Neill, Christine M. Wright, Bo-Ruei Chen, Wioletta Czaja, Rafael Contreras-Galindo

**Author notes:** To whom correspondence should be addressed to: Rafael Contreras-Galindo, Assistant Professor, Department of Genetics, University of Alabama at Birmingham, McCallum 627, 1918 University Blvd, Birmingham, Alabama, 35233, USA.

## Abstract

Systemic sclerosis (SSc) is a fibrotic autoimmune disease in which genomic sources of instability and their immunological consequences remain poorly defined. We show that bleomycin, a widely used SSc fibrosis model, induces DNA double-strand breaks (DSBs) at active centromeres. Similar centromeric damage signatures were observed in fibroblasts from patients with limited cutaneous SSc, consistent with prior observations. Quantification of α-satellite repeat content revealed dynamic changes in repeat abundance, consistent with deletions and insertions and incomplete restoration following damage. These breaks are repaired primarily ATM-dependent, RAD51-associated homologous recombination, but repair remains incomplete. Incomplete repair is associated with altered kinetochore assembly, chromosome missegregation, and increased formation of micronuclei and cytoplasmic chromatin enriched in centromere proteins. These fragments escape via nuclear envelope rupture and show spatial colocalization with MHC class II molecules. Together, these findings establish bleomycin-induced centromere damage as a tractable model to study active-centromere instability, its incomplete repair, and the resulting chromatin mislocalization in fibroblasts, with features relevant to systemic sclerosis.

## Introduction

Systemic sclerosis (SSc), or scleroderma, is a chronic autoimmune disease characterized by vascular injury, immune dysregulation, and progressive fibrosis of the skin and internal organs [1–4]. A defining clinical feature of limited cutaneous SSc (lcSSc) is the presence of anti-centromere antibodies (ACAs), directed against centromeric proteins such as CENP-A and CENP-B [5,6]. Yet the origin of these autoantigens remains unexplained. While recent progress has clarified immune pathways that sustain fibrosis [7–11], far less is known about how centromere fragility contributes to chromosomal instability and potential antigen exposure [2,12].

Human centromeres are composed of ∼171-bp α-satellite monomers arranged into higher-order repeat (HOR) arrays. These AT-rich repeats, of which only a subset harbor CENP-B boxes, are packaged into specialized chromatin marked by CENP-A [13,14]. Array size and organization vary extensively across individuals, with up to 30-fold differences recently revealed by near-complete genome assemblies [15]. While centromeres are essential for accurate chromosome segregation, their repetitive nature, secondary structure features and chromatin context make them prone to replication stress and DNA damage [16–20]. Damage within α-satellites disrupts kinetochore assembly and drives chromosomal instability (CIN), manifested by aneuploidy and micronucleus formation [17,21–23]. Mis-segregated chromosomes frequently form micronuclei that undergo rupture, releasing DNA into the cytoplasm. Cytosolic DNA can activate innate immune sensors such as cGAS, inducing interferon responses [2,24–26], and may enable interactions between nuclear-derived material and antigen-processing pathways [27–30]. These observations suggest a potential link between genome instability and immune activation, although direct connections to centromere-derived antigen presentation remain to be established.

Bleomycin (BLM), a chemotherapeutic antibiotic and widely used experimental fibrosis inducer, generates reactive oxygen species that produce both single- and double-strand DNA breaks (DSBs) [31,32]. Early studies demonstrated preferential BLM cleavage within repetitive DNA, including α-satellite sequences [33–36]. In vivo, BLM administration recapitulates key features of SSc, including dermal fibrosis, fibroblast activation, and extracellular matrix accumulation [37–39]. These properties make BLM a useful model to examine how genotoxic stress affects centromeric DNA and genome stability in fibroblasts and in vivo [2,40]. However, the repair dynamics of centromeric DSBs remain poorly understood. DSBs can be resolved by non-homologous end joining (NHEJ) or homologous recombination (HR), but in repetitive DNA, misalignment during HR can lead to deletions or insertions [41–43]. While CENP-A accumulation has been reported at induced damage sites [44,45], it is unclear whether this reflects recruitment of repair factors or the intrinsic fragility of centromeric chromatin.

Here, we use BLM-induced DNA damage in fibroblasts and a mouse skin fibrosis model to investigate centromere fragility and repair dynamics. Through α-satellite qPCR, immunofluorescence mapping, ATM inhibition, and patient fibroblast analyses, we demonstrate that BLM induces DSBs at active centromeres. These lesions are repaired primarily by RAD51-mediated HR but remain incompletely resolved, resulting in DNA loss, kinetochore disruption, missegregation, and micronucleus formation. Damaged centromere chromatin escapes into the cytoplasm through nuclear envelope rupture, where it shows spatial colocalization with HLA-DRB1. Together, these findings define a framework for studying active-centromere instability, its incomplete repair, and the resulting chromatin mislocalization in fibroblasts, with features relevant to systemic sclerosis.

## Results

### Centromere instability in BLM-induced scleroderma fibrosis models

To test whether BLM induces centromere instability in a fibrosis-relevant context, we first used the intradermal BLM mouse skin model, which reproduces dermal thickening, collagen deposition, and fibroblast activation. Mice received BLM injections every other day for 14 days. By day 14, fibrotic skin showed marked dermal thickness, erythema, and hydroxyproline compared with vehicle controls (Fig. 1A–D). Quantitative PCR (qPCR) of centromeric repeats revealed reduced relative abundance of centromeric repeats, with decreases of ∼50% for MaSat, ∼60% for MiSat, and ∼70% for Ymin in fibrotic tissue (Fig. 1E).

**Fig. 1.**
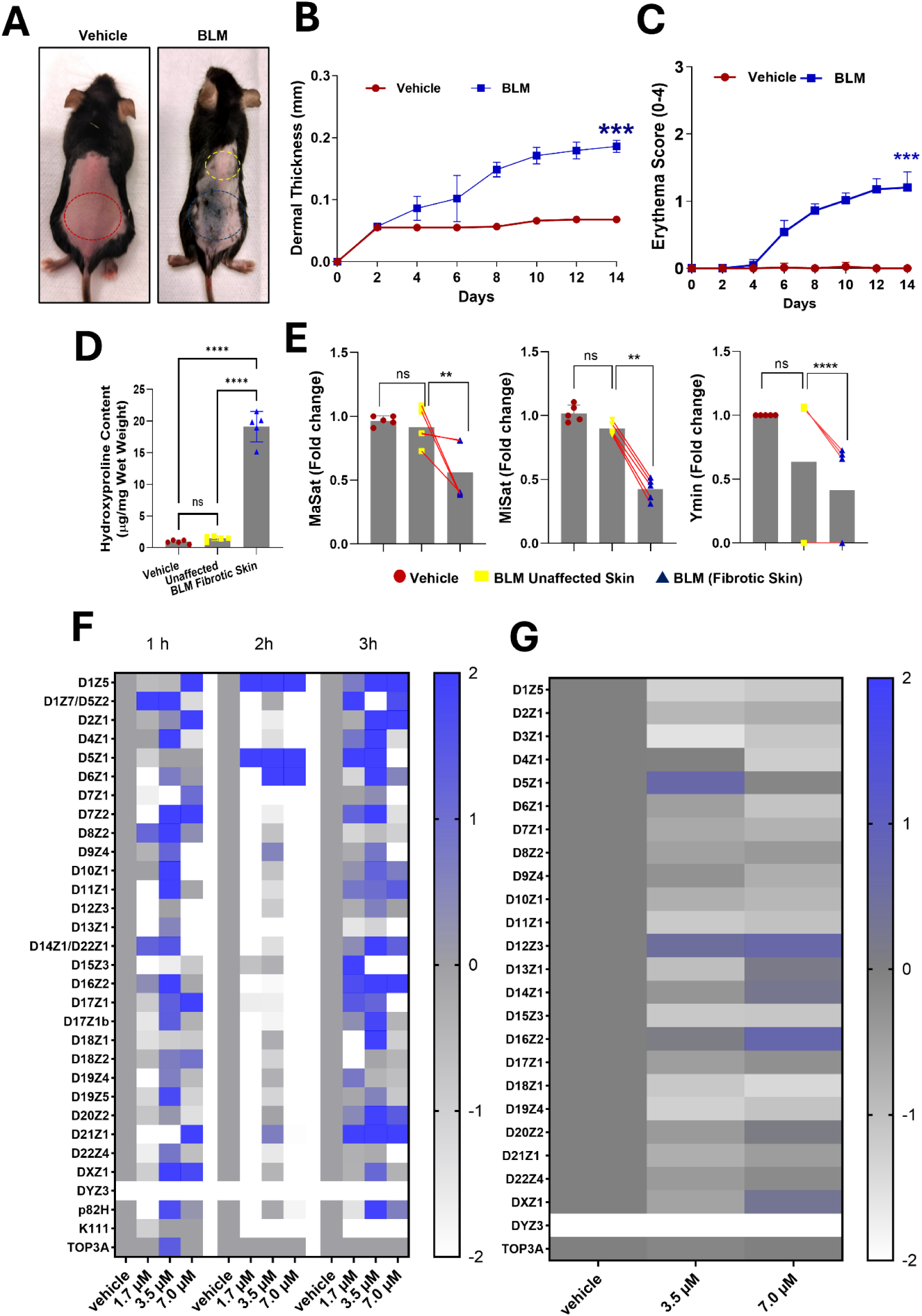
Bleomycin induces centromeric α-satellite alterations in mouse skin and human fibroblasts. (A) Representative dorsal skin images of mice after 14 days of vehicle (saline) or bleomycin (BLM) treatment (n = 5 per group). Red, vehicle-control skin; yellow, adjacent unaffected skin; blue, fibrotic lesion. (B–C) Quantification of dermal thickness (B) and erythema scores (C) in vehicle- and BLM-treated mice. (D) Hydroxyproline content measured by colorimetric assay in vehicle-control, adjacent unaffected, and fibrotic skin from BLM-treated mice. (E) qPCR analysis of mouse centromeric satellite repeats (MaSat, MiSat, Ymin) normalized to 18S. Red lines connect paired samples from adjacent unaffected and fibrotic skin within the same animal. (F–G) Heatmaps showing log₂ fold change in human α-satellite monomers in CHON-002 fibroblasts following BLM treatment. (F) Cells treated with BLM for 1–3 h. (G) Cells treated with BLM for 3 h followed by 24 h recovery. Color scale: white, loss; blue, gain; grey, stable. DYZ3 was not detected in CHON-002 cells (female origin). TOP3A served as a non-centromeric reference. Data are presented as mean ± SD from three independent experiments. Representative heatmaps are shown. Statistical significance was determined by one-way ANOVA with appropriate post hoc testing: \*\**p* < 0.01; \*\*\**p* < 0.001; \*\*\*\**p* < 0.0001; ns, not significant.

In human fibroblasts (CHON-002), BLM exposure (1.7–7.0 µM, 1–3 h) caused a rapid reduction in α-satellite DNA at 2 h, followed by partial recovery at 3 h, suggesting dynamic changes over time (Fig. 1F). At higher doses (7.0 µM), alterations persisted. After 24 h recovery, heatmap analysis showed sustained copy number alterations (predominantly losses, with occasional gains) across multiple α-satellite arrays, including ∼50% deletions at several centromere arrays (D1Z5, D6Z1, D15Z3, D18Z1) (Fig. 1G). BJ-5ta fibroblasts displayed a similar pattern but the extent of centromere alterations was more severe, with deletions and insertions occurring at comparable frequency, and persistent losses at D5Z1, D7Z1, D9Z4, and D17Z1 (Supplementary Fig. 1A–B).

MTT assays confirmed ∼75–80% viability at 3.5–7.0 µM (Supplementary Fig. 2), indicating that centromere copy number changes are not attributable to overt cytotoxicity under these conditions.

### BLM-induced double-strand breaks at active centromeres

We next asked whether BLM preferentially induces DSBs at centromeres marked by CENP-A. Here, we define active centromeres as CENP-A–containing chromatin domains, which mark functional kinetochore-associated centromeres. CHON-002 fibroblasts treated with 3.5 or 7.0 µM BLM for 3 h showed increased γ-H2AX signal compared with untreated controls, consistent with widespread DSB induction (Fig. 2A). Similar results were obtained in BJ-5ta fibroblasts (Supplementary Fig. 3A), and western blotting confirmed increased γ-H2AX levels following BLM exposure (Supplementary Fig. 3B).

**Fig. 2.**
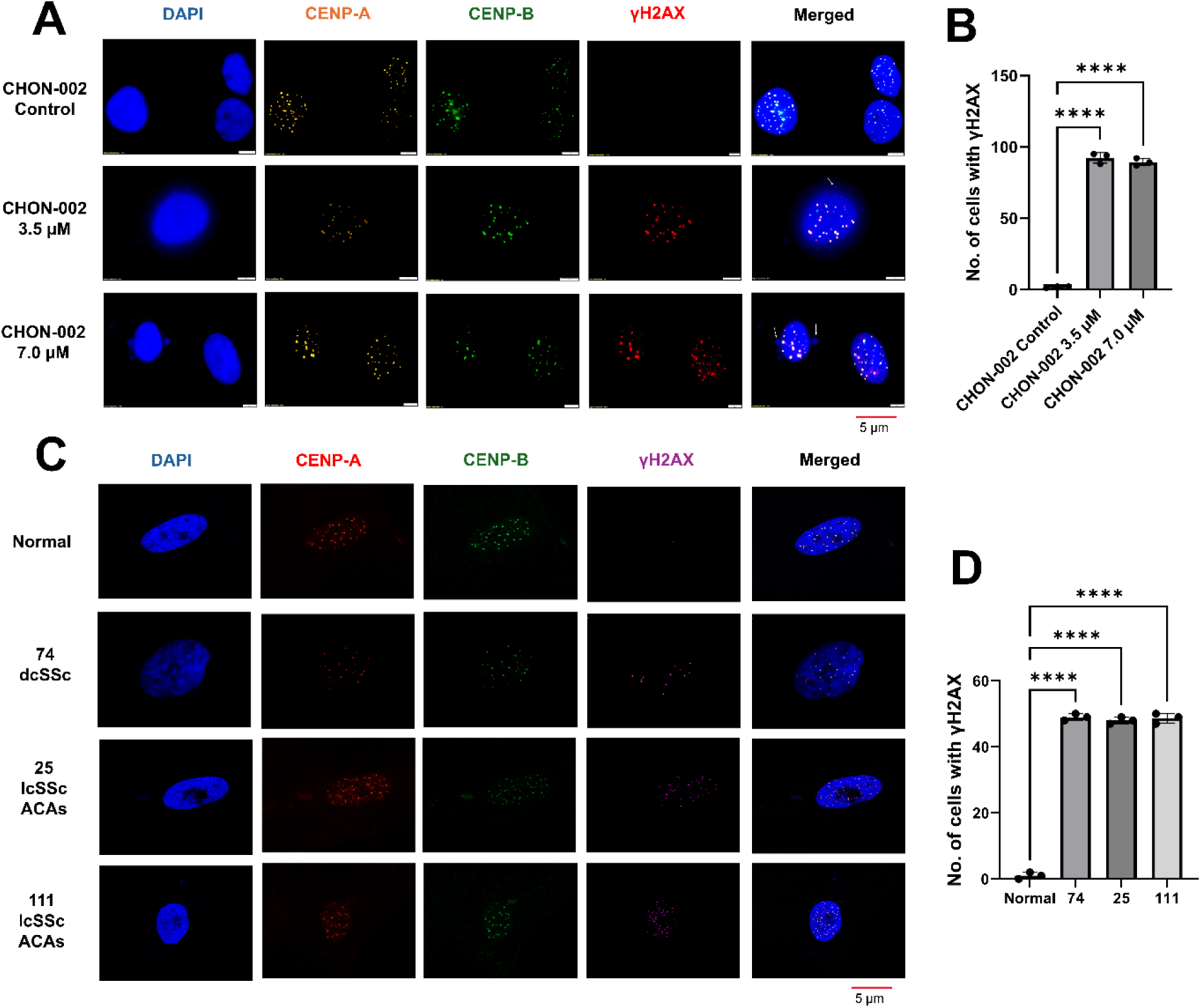
Bleomycin induces DNA damage at centromere-associated regions in fibroblasts. (A) Immunofluorescence analysis of CHON-002 fibroblasts treated with bleomycin (BLM; 3.5 or 7.0 µM, 3 h) and stained for CENP-A (orange), CENP-B (green), γ-H2AX (red), and DAPI (blue, nuclei). White arrows indicate micronuclei. (B) Quantification of γ-H2AX-positive cells (n = 100 cells). (C) Immunofluorescence analysis of primary dermal fibroblasts from healthy controls and SSc patients stained for CENP-A (green), γ-H2AX (purple), and DAPI (blue). (D) Quantification of γ-H2AX-positive cells (n = 50 cells). CENP-A and CENP-B signals were detected in all nuclei across conditions. Data represent mean ± SD from three independent counts. Statistical significance was determined using unpaired t-test or one-way ANOVA: \**p* < 0.05; \*\**p* < 0.01; \*\*\**p* < 0.001; ns, not significant. Quantitative correlation analysis is shown in Fig. S3.

Colocalization analysis revealed that γ-H2AX foci frequently overlapped with centromeric markers. While CENP-A and CENP-B signals remained tightly correlated (r > 0.85) under all conditions, γ-H2AX showed higher pixel intensity correlation with CENP-A (r = 0.51–0.74) than with CENP-B (r = 0.31–0.50), consistent with increased association of γH2AX with CENP-A–marked regions (Supplementary Fig. 3C–D).

To assess disease relevance, we examined fibroblasts from three SSc patients. γ-H2AX signal was detected in all samples. In fibroblasts from two lcSSc patients, γ-H2AX showed overlap with CENP-A (Pearson’s r = 0.29–0.42), with lower overlap observed with CENP-B (r = 0.09–0.19). In contrast, fibroblasts from one dcSSc patient displayed γ-H2AX signal with minimal overlap with CENP-A (Fig. 2B, Supplementary Fig. 3E).

Together with the BLM experiments, these findings suggest that acute fibroblast models and patient-derived fibroblasts show distinct patterns of γ-H2AX association with centromeric markers. In lcSSc fibroblasts, a subset of cells showed γ-H2AX association with CENP-A, whereas dcSSc fibroblasts displayed γ-H2AX signal with limited overlap with CENP-A. The small number of samples precludes definitive conclusions regarding disease subset–specific mechanisms.

### BLM-induced centromeric breaks are associated with RAD51-marked homologous recombination

To determine the DNA repair pathways engaged at centromeric DSBs, CHON-002 fibroblasts were treated with 3.5 or 7.0 µM BLM and immunostained for γ-H2AX (DSB marker), RAD51 (HR), and KU70 (NHEJ). Untreated cells showed minimal nuclear staining (Fig. 3A). BLM exposure induced γ-H2AX foci with associated RAD51 signal, while KU70 signal showed lower overlap with γ-H2AX.

**Fig. 3.**
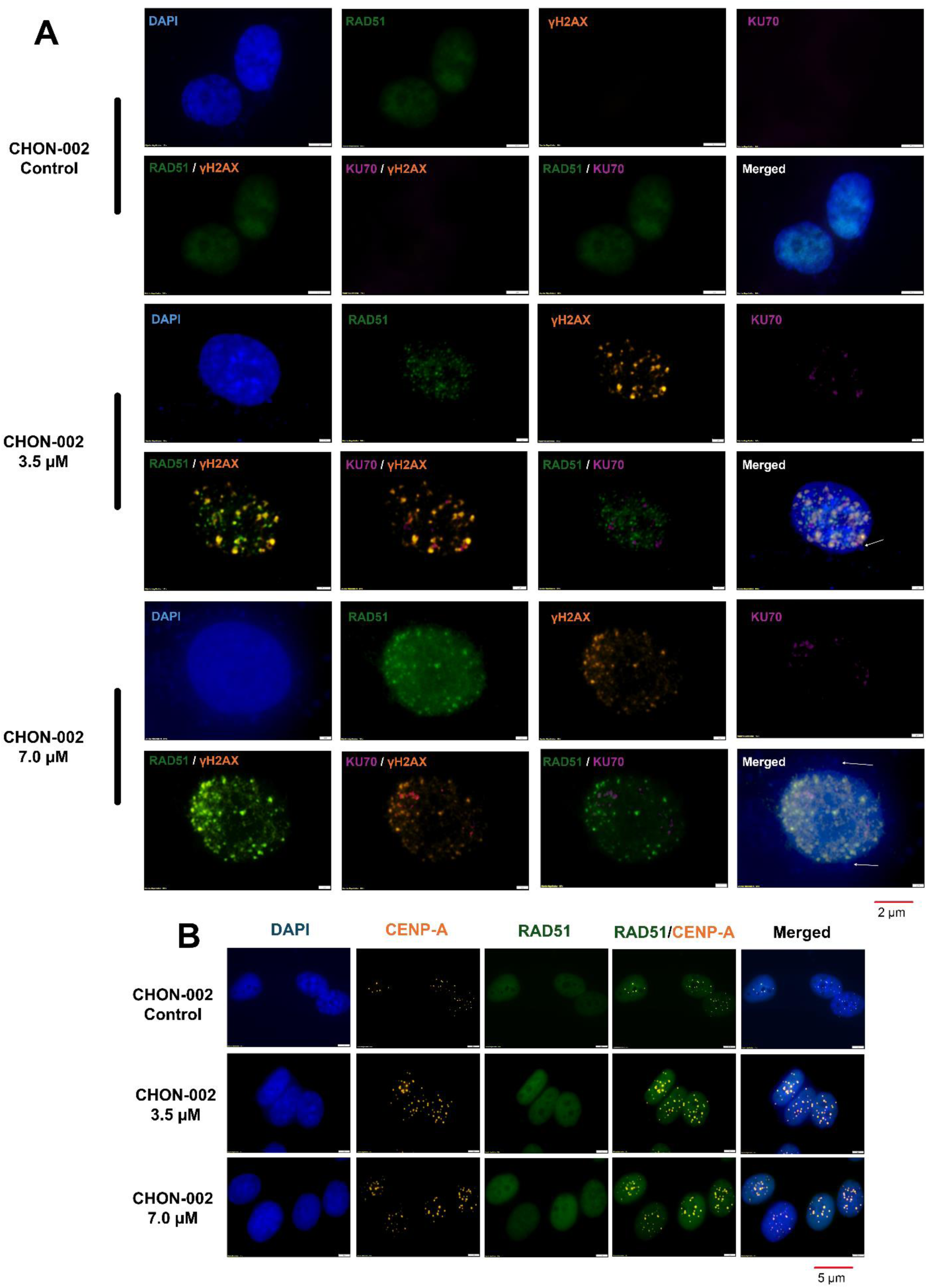
Association of homologous recombination factors to centromeric regions following bleomycin treatment. CHON-002 fibroblasts were treated with bleomycin (BLM; 3.5 or 7.0 µM) for 3 h followed by 24 h recovery, unless otherwise indicated. (A) Immunofluorescence analysis of RAD51 (homologous recombination; green), γ-H2AX (DNA double-strand breaks; orange), KU70 (non-homologous end joining; purple), and DAPI (blue, nuclei). White arrows indicate micronuclei. (B) Immunofluorescence analysis of CENP-A (orange) and RAD51 (green) following 3 h BLM treatment (no recovery). RAD51 signal is shown in relation to centromeric regions marked by CENP-A. Quantitative correlation analysis is shown in Fig. S4.

Quantitative analysis showed higher association of RAD51 with γ-H2AX compared to KU70. At 3.5 µM BLM, RAD51–γ-H2AX correlation was high (r = 0.875), compared with lower values for KU70–γ-H2AX (r = 0.314). At 7.0 µM, RAD51–γ-H2AX correlation further increased (r = 0.919), whereas KU70 associations remained low (r = 0.276) (Supplementary Fig. 4A). Gene expression analysis showed that RAD51 mRNA increased in a dose-dependent manner (****p < 0.0001), while KU70 showed only a modest but significant rise (**p < 0.001, Supplementary Fig. 4B). Kinetic analyses revealed temporal differences in repair factor localization. At 3 h post-treatment, RAD51 staining was diffuse, but by 24 h distinct RAD51 foci colocalized with γ-H2AX (Fig. 3A–B).

### BLM activates PARP-dependent single-strand break and replication stress signaling

To assess the single-strand break (SSB) and replication stress response, we monitored poly(ADP-ribose) (PAR) polymer accumulation, a rapid marker of PARP activation [41]. In untreated CHON-002 fibroblasts, PAR immunofluorescence was minimal or absent (Supplementary Fig. 4C). Exposure to 7.0 µM BLM induced increased nuclear PAR signal in ∼20% of cells, where the signal was pan-nuclear and diffuse, consistent with PARP activation following DNA damage. The remaining cells showed low or undetectable PAR signal, reflecting the rapid turnover of PAR polymers. Quantification confirmed a significant increase in nuclear PAR intensity compared with untreated controls (****p < 0.0001, n = 50 cells) (Supplementary Fig. 4D). Although PAR staining indicated an SSB response, the BLM exposure was associated with widespread γ-H2AX accumulation, consistent with DSB formation, as reflected by widespread γ-H2AX accumulation.

### ATM-mediated phosphorylation is required for γ-H2AX and RAD51 recruitment to centromeric DSBs

ATM kinase phosphorylates H2AX to generate γ-H2AX at sites of DNA damage. To test whether ATM mediates BLM-induced centromeric signaling, CHON-002 fibroblasts were treated with 3.5 or 7.0 µM BLM for 3 h, with or without the ATM inhibitor KU-55933 (35 µM, 1 h pretreatment). BLM was associated with dose-dependent increases in γ-H2AX signal, with partial overlap with the centromeric marker CENP-A (Fig. 4A). ATM inhibition decreased γ-H2AX intensity at both doses, while CENP-A localization remained intact (Fig. 4A-C). Furthermore, ATM inhibition reduced RAD51 puncta accumulation (Fig. 4D).

**Fig. 4.**
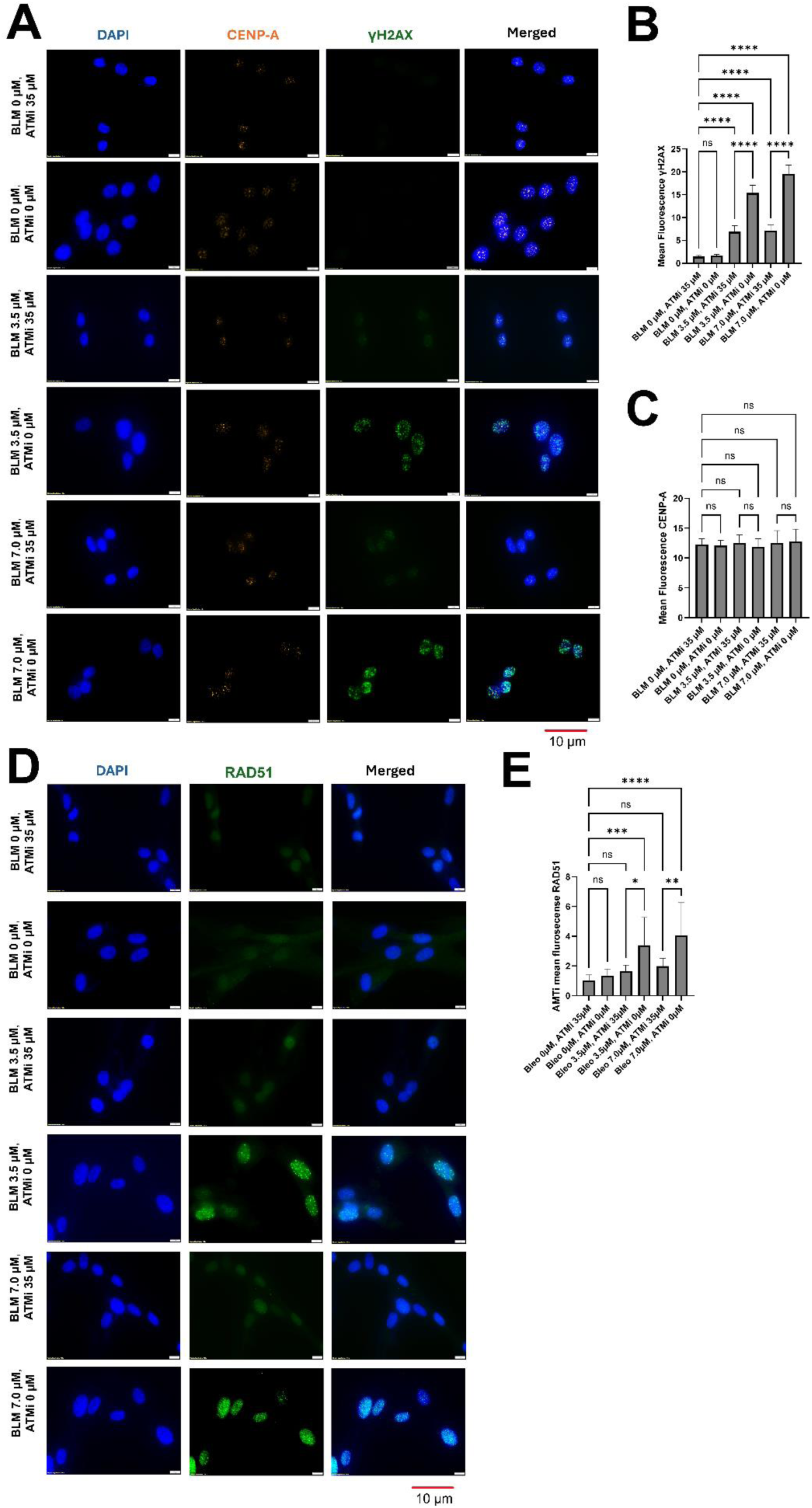
ATM inhibition modulates γ-H2AX and RAD51 signals following bleomycin treatment. CHON-002 fibroblasts were treated with bleomycin (BLM; 3.5 or 7.0 µM) for 3 h, with or without 1 h pretreatment with the ATM inhibitor KU-55933 (35 µM). (A) Immunofluorescence analysis of DAPI (blue, nuclei), CENP-A (orange, centromeres), and γ-H2AX (green, DNA double-strand breaks). γ-H2AX signal is shown relative to centromeric regions marked by CENP-A. Scale bar, 10 µm. (B) Quantification of mean nuclear γ-H2AX intensity across conditions (n = 50 cells per condition). (C) Quantification of mean nuclear CENP-A intensity across conditions. (D) Immunofluorescence analysis of DAPI (blue, nuclei) and RAD51 (green, homologous recombination). Scale bar, 10 µm. (E) Quantification of mean nuclear RAD51 intensity across conditions (n = 50 cells per condition). Data are presented as mean ± SD. Statistical significance was determined by one-way ANOVA with appropriate post hoc testing: \**p* < 0.05; \*\**p* < 0.01; \*\*\**p* < 0.001; \*\*\*\**p* < 0.0001; ns, not significant.

Quantitative analysis confirmed that γ-H2AX and RAD51 induction by BLM was significantly suppressed by ATM inhibition (\*\*\**p* < 0.001, \*\*\*\**p* < 0.0001, one-way ANOVA) (Fig. 4B,E). In contrast, CENP-A fluorescence levels were unaffected (Fig. 4C, ns). These findings indicate that ATM contributes to γ-H2AX signaling and RAD51 association under these conditions at centromeric breaks.

### BLM-induced DNA damage triggers micronuclei formation and centromere chromatin release

DAPI staining revealed increased frequency of micronuclei in CHON-002 and BJ-5ta fibroblasts treated with 3.5 or 7.0 µM BLM, whereas micronuclei were infrequent in controls (Fig. 5A and Supplementary Fig. 5A). Many micronuclei contained centromeric signals (Fig. 5A and Supplementary Fig. 5A), consistent with observations in SSc fibroblasts [2]. However, a subset of micronuclei contained CENP-B but lacked detectable CENP-A signal (Fig. 5A–D, Supplementary Fig. 5A–E), and this pattern was consistently observed across conditions, supporting disruption of centromere organization as previously described in SSc fibroblasts [2]. In untreated cells, CENP-A and CENP-B were localized exclusively to nuclear centromeres, consistent with patterns in healthy skin fibroblasts [2]. After BLM exposure, centromere-associated signals were detected in the cytoplasm. At 7.0 µM, ∼12% of cells displayed cytoplasmic centromere-associated structures (\*\*\*\**p* < 0.0001, n = 3) (Fig. 5E), a finding confirmed in 15% of BJ-5ta fibroblasts (Supplementary Fig. 5A, F, \*\*\*\**p* < 0.0001, n = 3), reinforcing that BLM promotes the presence of cytoplasmic centromere-associated structures.

**Fig. 5.**
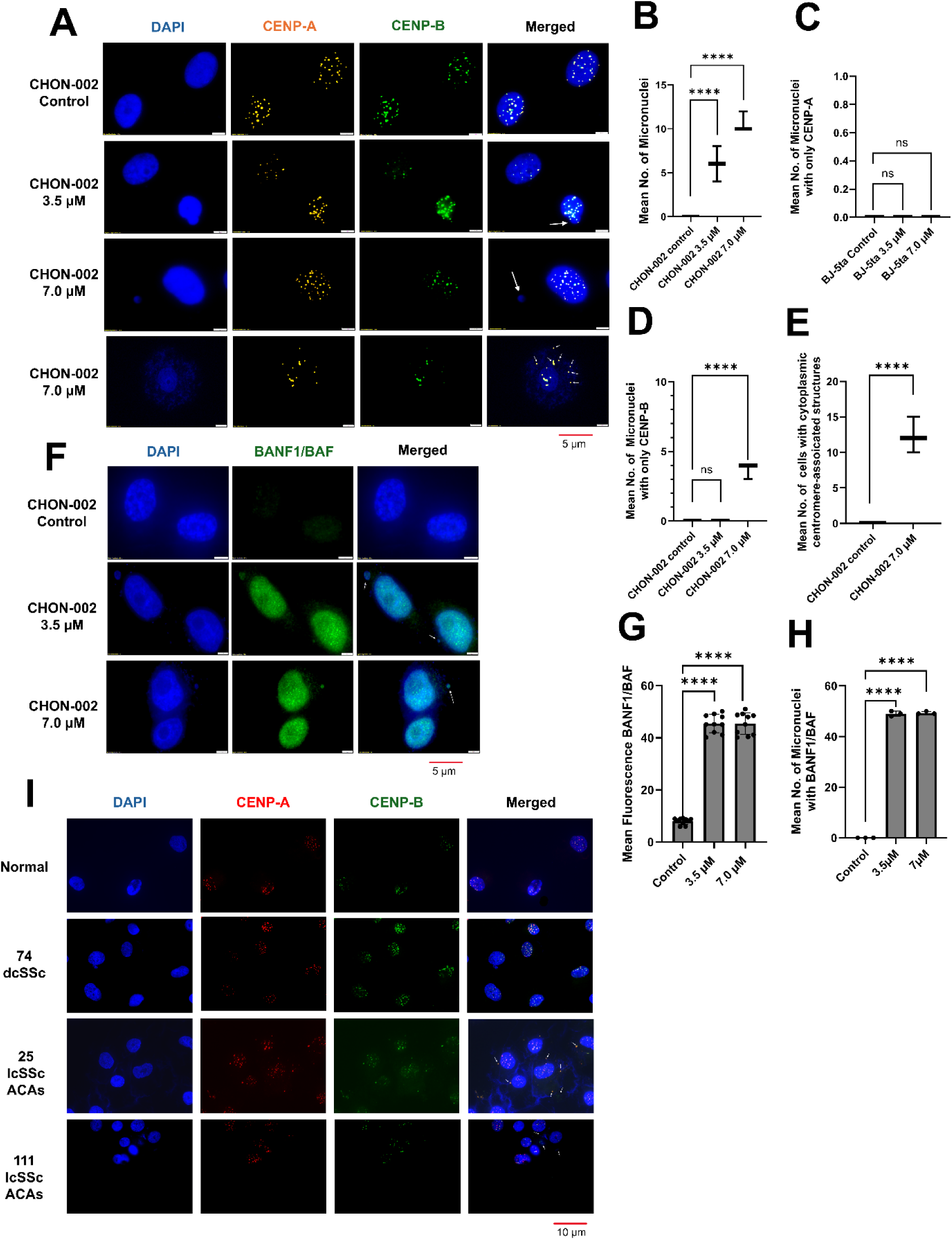
Bleomycin treatment is associated with micronuclei formation, cytoplasmic centromeric chromatin, and altered BANF1 localization. CHON-002 fibroblasts were treated with bleomycin (BLM; 3.5 or 7.0 µM) for 3 h followed by 24 h recovery, unless otherwise indicated. (A) Immunofluorescence analysis of CENP-A (orange), CENP-B (green), and DAPI (blue, nuclei). White arrows indicate micronuclei; small white arrows indicate cytoplasmic centromere-associated structures. B–E) Quantification of (B) mean number of micronuclei per 100 cells, (C) mean number of micronuclei positive for CENP-A, (D) mean number of micronuclei positive for CENP-B, and (E) mean number of cells containing cytoplasmic centromere-associated structures (per 100 cells). Quantifications are based on 100 cells per condition. Data represent mean ± SD from three independent experiments. (F) Immunofluorescence analysis of BANF1 (green) and DAPI (blue). BANF1 signal is shown in nuclei and micronuclei (white arrows). (G) Quantification of mean BANF1 fluorescence intensity across conditions (n = 50 cells per condition). (H) Quantification of micronuclei positive for BANF1, expressed as the percentage of micronuclei positive for BANF1 (n = 200 micronuclei). (I) Immunofluorescence analysis of primary dermal fibroblasts from healthy controls and systemic sclerosis (SSc) patients stained for CENP-A (red), CENP-B (green), and DAPI (blue). White arrows indicate cytoplasmic centromere-associated structures in two lcSSc patients (25 and 111), but not observed in a dcSSc patient (74). Data are presented as mean ± SD. Statistical significance was determined by one-way ANOVA with appropriate post hoc testing: \*\*\*\**p* < 0.0001; ns, not significant.

To test whether nuclear envelope rupture mediates this release, we examined BANF1, a marker of nuclear envelope perturbation. BANF1 was localized to nuclei in control cells but strongly accumulated at rupture sites on nuclei and micronuclei in BLM-treated fibroblasts (white arrows, Fig. 5F). Quantification confirmed significantly elevated BANF1 intensity (****p < 0.0001, n = 50) (Fig. 5G) and an increased percentage of micronuclei positive for BANF1 (Fig. 5H).

Patient fibroblasts showed similar phenotypes [2]. Consistently, two lcSSc fibroblasts (patients 25 and 111) frequently displayed cytoplasmic centromeric chromatin, whereas cells from one dcSSc patient rarely did (Fig. 5I). This pattern is consistent with the higher prevalence of ACAs among lcSSc patients. Together with BANF1 redistribution, these findings are consistent with altered nuclear envelope integrity contributing to cytoplasmic chromatin presence in lcSSc fibroblasts.

### Centromere-associated signals show spatial proximity to MHC class II components

BLM-induced appearance of centromere-associated chromatin in the cytoplasm was observed in treated fibroblasts, consistent with prior reports in lcSSc [2]. To examine the spatial relationship between centromere-associated signals and MHC class II components, we performed IF for CENP-B and HLA-DRB1. Treated fibroblasts showed partial spatial proximity in the cytoplasm between CENP-B and HLA-DRB1 (Fig. 6). This overlap was observed in cells that exhibited cytoplasmic centromere-associated signals. Fibroblasts from ACA-positive lcSSc patients showed similar patterns of spatial overlap, whereas cells lacking cytoplasmic centromere-associated signals showed minimal overlap. These observations indicate that cytoplasmic CENP-B signals are found in proximity to MHC class II components under these conditions.

**Fig. 6.**
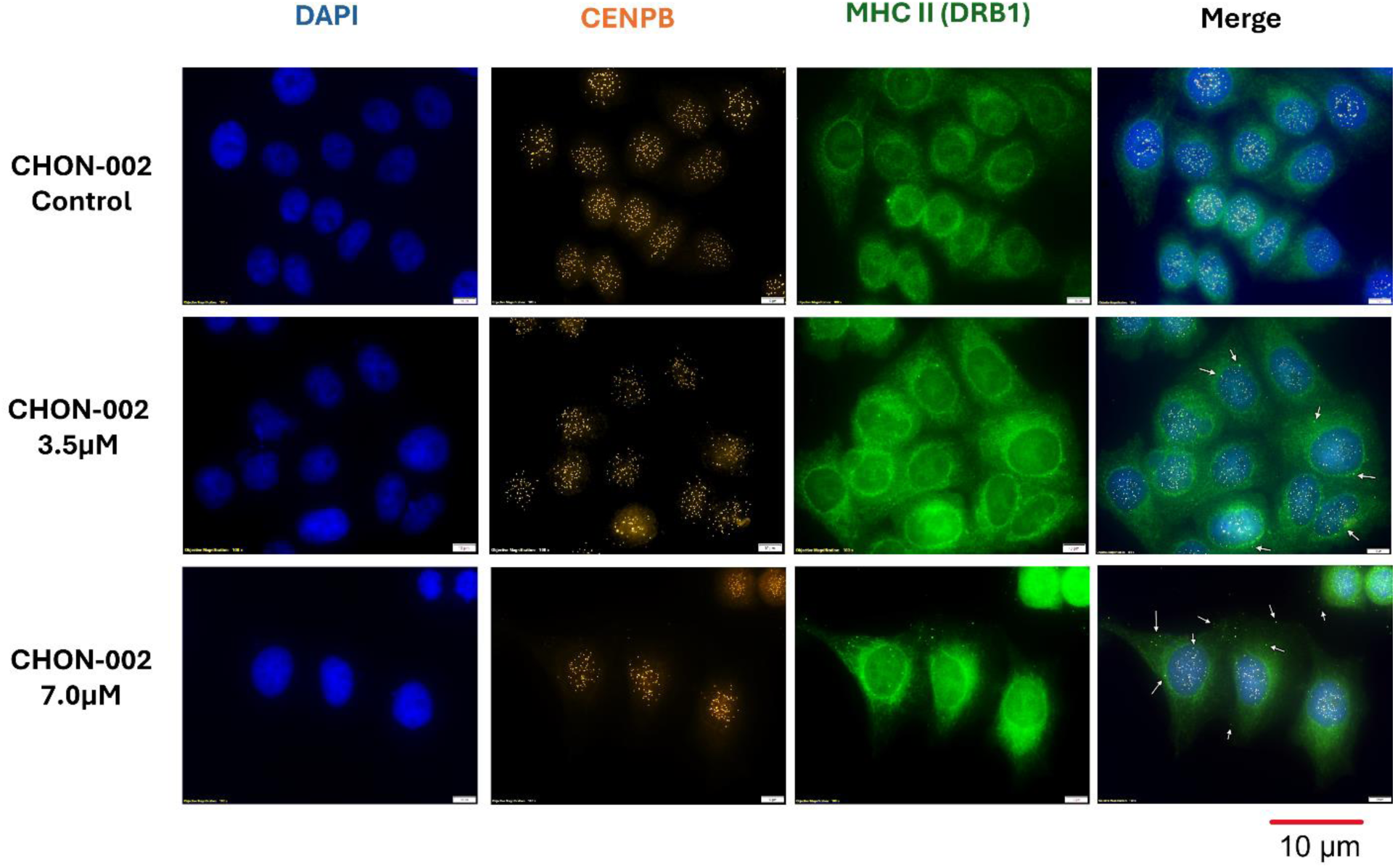
Partial spatial overlap of cytoplasmic CENP-B and MHC class II (HLA-DRB1) following bleomycin treatment. CHON-002 fibroblasts were left untreated or treated with bleomycin (BLM; 3.5 or 7.0 µM) for 3 h followed by 24 h recovery. Immunofluorescence analysis of CENP-B (orange), MHC class II (HLA-DRB1; green), and DAPI (blue, nuclei). White arrows indicate cytoplasmic regions where CENP-B and HLA-DRB1 signals overlap.

## Discussion

This study identifies centromeric DNA as a target of genotoxic stress in fibroblasts. We demonstrate that bleomycin (BLM)-induced breaks within centromeric α-satellite arrays are signaled through ATM-dependent γ-H2AX and repaired by RAD51-mediated homologous recombination (HR), yet remain incompletely resolved, resulting in deletions and insertions within centromeric repeats. One consequence of this incomplete repair is the appearance of centromere-associated chromatin into the cytoplasm, where it shows spatial proximity with MHC class II molecules. Similar features observed in patient-derived fibroblasts and in the BLM-induced skin fibrosis model support the relevance of these processes to disease-associated cellular phenotypes. These findings position centromeres as active participants in genome stress responses rather than solely structural elements.

Centromeric repeats are intrinsically vulnerable due to their repetitive sequence composition, replication challenges, and specialized chromatin organization. Prior studies have shown that centromeres accumulate spontaneous lesions during both proliferation and quiescence, with repair occurring predominantly through RAD51-mediated HR [18,20,46]. In this study, BLM exposure amplified this baseline fragility, generating deletions and insertions within α-satellite arrays. The magnitude of these alterations differed between fibroblast lines, suggesting that chromatin state or repair capacity may influence centromeric vulnerability. Consistent with these observations, the BLM-induced skin fibrosis model recapitulated reductions in centromeric satellite content in vivo [32–35]. Together, these findings support centromeric DNA as a sensitive substrate for genotoxic stress across experimental systems.

Colocalization of γ-H2AX with CENP-A and suppression of this signal by ATM inhibition demonstrate that centromeric lesions engage canonical ATM-dependent DNA damage signaling. Preferential recruitment of RAD51, with limited KU70 involvement, indicates that HR is the dominant repair pathway at these sites. However, HR did not fully restore centromeric array integrity, consistent with prior observations that repair within repetitive DNA can be error-prone due to template misalignment or replication-associated stress [20,46,47]. The presence of micronuclei containing CENP-B in the absence of CENP-A further suggests that centromere identity and structural integrity can become disrupted under these conditions, consistent with prior observations in SSc fibroblasts [2].

Functionally, centromere instability was associated with chromosome missegregation, micronucleus formation, and nuclear envelope rupture, marked by BANF1 redistribution and cytoplasmic appearance of centromeric chromatin. These phenotypes were observed in a subset of BLM-treated fibroblasts and were consistent with features previously described in systemic sclerosis (SSc) fibroblasts, particularly in limited cutaneous SSc [2]. Cytoplasmic CENP-B showed spatial proximity with HLA-DRB1, suggesting that mislocalized centromeric chromatin may come into proximity with antigen-processing compartments. These observations are consistent with prior work linking cytosolic DNA to innate immune signaling pathways, including cGAS–STING, and suggest that centromeric DNA may contribute to the pool of cytosolic DNA generated during genome instability [2,5,48–51].

Differences observed between fibroblast lines, including lcSSc and dcSSc samples, suggest variability in centromere damage responses and repair dynamics. However, the limited number of patient-derived lines analyzed here precludes definitive conclusions regarding disease subset–specific mechanisms.

This study has limitations. Fibroblast cultures and the BLM model do not fully capture the complexity of stromal-immune interactions in vivo, and direct demonstration of centromeric peptide processing, antigen presentation, and downstream immune activation remains to be established. We note that colocalization analyses do not establish direct molecular interactions or antigen presentation.

In summary, these findings define a framework for studying active-centromere instability, its incomplete repair, and the resulting centromere-associated chromatin mislocalization in fibroblasts. This system provides a tractable platform to investigate how damage to repetitive genomic regions contributes to genome instability and may influence cellular pathways relevant to fibrosis and autoimmunity. Future studies will be required to determine how these processes intersect with immune signaling and disease progression [52–55].

## Methods

### Reagents

Bleomycin sulfate (Cayman Chemical, Cat. No. 13877, Lot #0701763-49) was used in all experiments for consistency across experiments. All other reagents were of analytical grade and obtained from commercial suppliers unless otherwise noted.

### Cell culture and treatments

hTERT-immortalized human fibroblasts (CHON-002, ATCC CRL-2847; BJ-5ta, ATCC CRL-4001) were cultured in Dulbecco’s Modified Eagle Medium (DMEM; Gibco) supplemented with 10% fetal bovine serum (FBS; Gibco) and 1% penicillin–streptomycin (Gibco). Cells were maintained at 37 °C in a humidified incubator with 5% CO₂ and routinely tested for mycoplasma contamination.

For acute bleomycin (BLM) exposure, fibroblasts were seeded at 2-3 × 10⁵ cells per well in 6-well plates and grown to 60–80% confluence. Cells were treated with 0, 3.5, or 7 µM BLM for 3 h. For recovery assays, cells were washed with phosphate-buffered saline (PBS) and incubated in drug-free medium for 24 h. For ATM inhibition, cells were pretreated with KU-55933 (35 µM, Selleck Cat. S1092) for 1 h prior to BLM treatment.

### Mouse BLM-induced skin fibrosis model

C57BL/6 mice (Jackson Laboratory; 20–25 g; 2 males, 3 females per group) were randomized to receive intradermal injections of BLM (1 U/mL, 100 µL) or vehicle (PBS) into shaved dorsal skin every other day for 14 days. On day 14, mice were anesthetized with isoflurane (1-2%) and euthanized by cervical dislocation. Skin samples were collected from: (1) PBS-injected control sites, (2) adjacent uninjected skin from the same animal, and (3) fibrotic BLM-injected lesions. Genomic DNA was isolated for qPCR analysis. All animal studies were approved by the University of Alabama at Birmingham Institutional Animal Care and Use Committee (protocol 23172).

### Human samples

Primary dermal fibroblasts were derived from patients with SSc under University of Michigan IRB HUM00065044, as previously described [2]. Cell lines were obtained from patients with limited cutaneous SSc (lcSSc) and diffuse cutaneous SSc (dcSSc), as well as from healthy controls.

### DNA and RNA isolation

Genomic DNA was extracted using the Monarch Genomic DNA Purification Kit (NEB 30102) with RNase A treatment. Total RNA was isolated using the Direct-zol RNA MicroPrep Kit (Zymo Research) with on-column DNase digestion. Concentrations were measured with a Qubit 4 Fluorometer (Thermo Fisher). DNA and RNA were stored at −80 °C until use.

### Centromere qPCR

Mouse centromeric arrays (MaSat, MiSat, and Ymin) were quantified by qPCR using validated primer sets [56]. Human α-satellite arrays (D1–D22, X, Y) were quantified using primer sets described previously [57]. Reactions were run on QuantStudio 3 (Applied Biosystems) with Radiant SYBR Green Master Mix (Alkali Scientific). Relative copy number was normalized to TOP3A (human) or 18S (mouse), unless otherwise indicated. Primer specificity was confirmed by melt-curve analysis.

### Western blotting

Cells were lysed in RIPA buffer (ChemCruz) supplemented with protease and phosphatase inhibitors (Thermo Fisher). Lysates (20–30 µg protein) were resolved by SDS–PAGE (10%) and transferred to PVDF membranes (Bio-Rad). Membranes were blocked in 5% milk/TBST, incubated overnight at 4 °C with primary antibodies (Supplementary Table 1), and probed with HRP-conjugated secondary antibodies (2 h, room temperature). Signals were developed using Clarity ECL substrate (Bio-Rad) and imaged with a ChemiDoc MP system. Band intensities were quantified using Image Lab (Bio-Rad) or ImageJ when applicable.

### Immunofluorescence microscopy

Cells were fixed in 4% paraformaldehyde (PFA), permeabilized with PBST (0.2% Triton X-100), and blocked in 2% BSA/PBST. Coverslips were incubated with primary antibodies (Supplementary Table 1) for 1 h at 37 °C, followed by Alexa Fluor–conjugated secondary antibodies (1:1000, 45 min, room temperature). Nuclei were counterstained with DAPI (ProLong Gold, Thermo Fisher).

For chromosome spreads, cells were arrested with 10 µg/mL colcemid for 16 h, incubated in 0.075 M KCl for 15 min, cytospun at 290 × g for 5 min, and fixed in 4% PFA prior to IF staining. Images were acquired on an Olympus BX73 microscope using identical exposure settings across conditions and processed uniformly using Fiji/ImageJ. Colocalization analyses were performed using pixel intensity correlations (Pearson’s r) with the Coloc2 plugin in Fiji/ImageJ using consistent thresholding parameters across conditions.

### Quantitative RT-PCR

mRNA expression of RAD51 and KU70 was measured using the Luna One-Step RT-qPCR Kit (NEB) with 50 ng RNA per reaction on a QuantStudio 3 instrument. GAPDH was used as a reference gene, and relative expression was calculated using the ΔΔCt method. Reactions were performed in technical triplicates. Primer sequences were obtained from published sources [58,59].

### Statistical analysis

All qPCR experiments were performed in three independent biological replicates with technical duplicates. Immunofluorescence quantifications were performed on at least 50–100 cells per condition from randomly selected fields, unless otherwise specified. Western blotting was performed in at least three independent replicates. Statistical significance was determined by Unpaired two-tailed Student’s t-test or one-way ANOVA with Tukey’s or Dunnett’s post hoc test, as appropriate (GraphPad Prism v9.3.1). A threshold of p < 0.05 was considered significant. Colocalization coefficients (Pearson’s r) were calculated from pixel intensity correlations using the Coloc2 plugin in Fiji/ImageJ with consistent thresholding parameters applied across all conditions.

## Data availability

The data generated in this study are provided in the Source Data file. Source data are provided with this paper.

## Acknowledgements

We thank Dr. Arko Sen for insightful discussions. The authors thank the National Scleroderma Foundation for supporting this project. R.C.-G. is the recipient of the Marta Marx Award for the Eradication of Scleroderma from the National Scleroderma Foundation.

## Author contributions

R.C-G. and A.I. conceived and designed the study. R.C-G., A.I., M.W. and C.W designed and performed the investigation. R.C-G acquired the funding. R.C-G., M.W., B.C., and W.C supervised the study. R.C-G., M.W., H.O., B.C., and W.C. wrote, reviewed and edited the manuscript.

## Additional information

Supplementary Information is available for this paper.

