## Supplementary Information for "Bleomycin induces active-centromere damage and cytoplasmic mislocalization of centromeric chromatin in fibroblasts"

**Centromere Instability Drives Chromosome Damage and Autoantigen Exposure in Systemic Sclerosis**
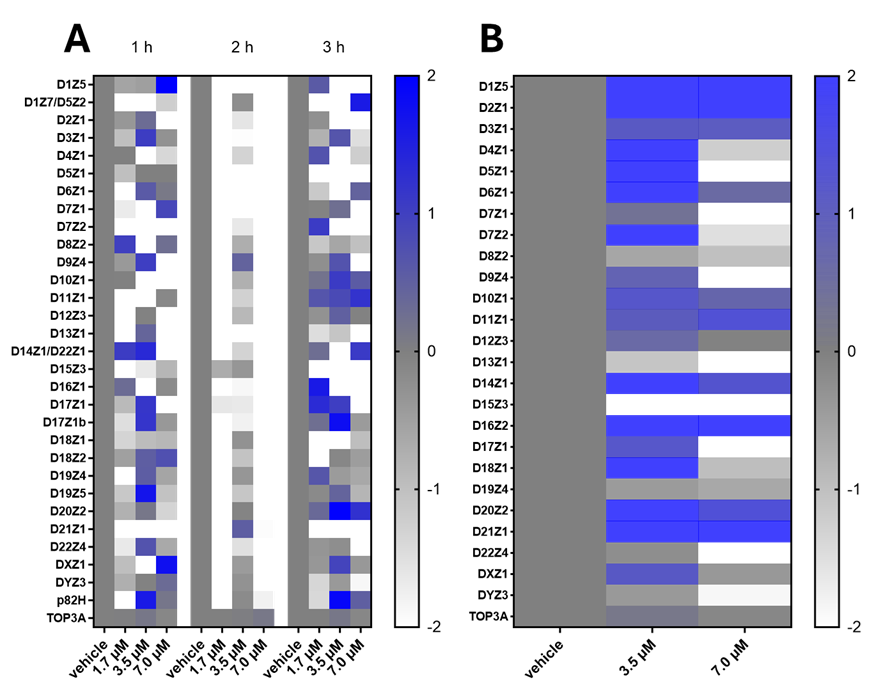

**Supplementary Fig. 1. Heatmap analysis of α-satellite abundance in BJ-5ta fibroblasts after BLM.** (A) Heatmaps showing log₂ fold-change in α-satellite monomers after treatment with 1.7, 3.5, or 7.0 µM BLM for 1–3 h. (B) Heatmap showing changes after 3 h BLM treatment followed by 24 h recovery. Color scale is the same as Fig. 1. Data represent biological duplicates from three independent experiments; representative heatmaps are shown.

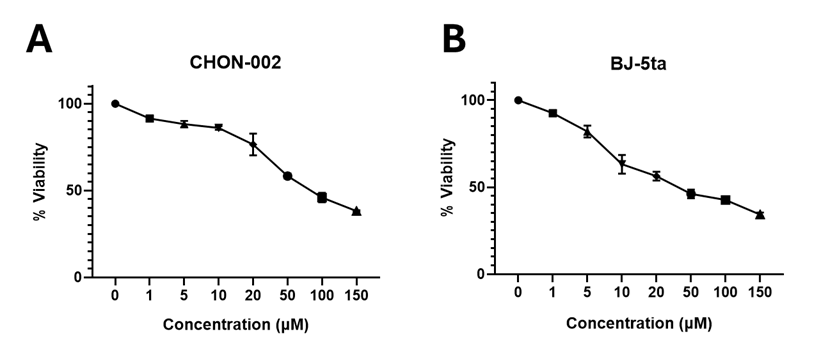

**Supplementary Fig. 2. Cytotoxicity of BLM in fibroblasts.** (A) CHON-002 and (B) BJ-5ta fibroblasts treated with 1–150 µM BLM for 3 h; viability was measured 24 h later by MTT assay. Cell viability is expressed relative to untreated controls (100%). Data represent mean ± SD from three independent experiments. Estimated LD₅₀ values were ~80 µM for CHON-002 and ~45 µM for BJ-5ta.

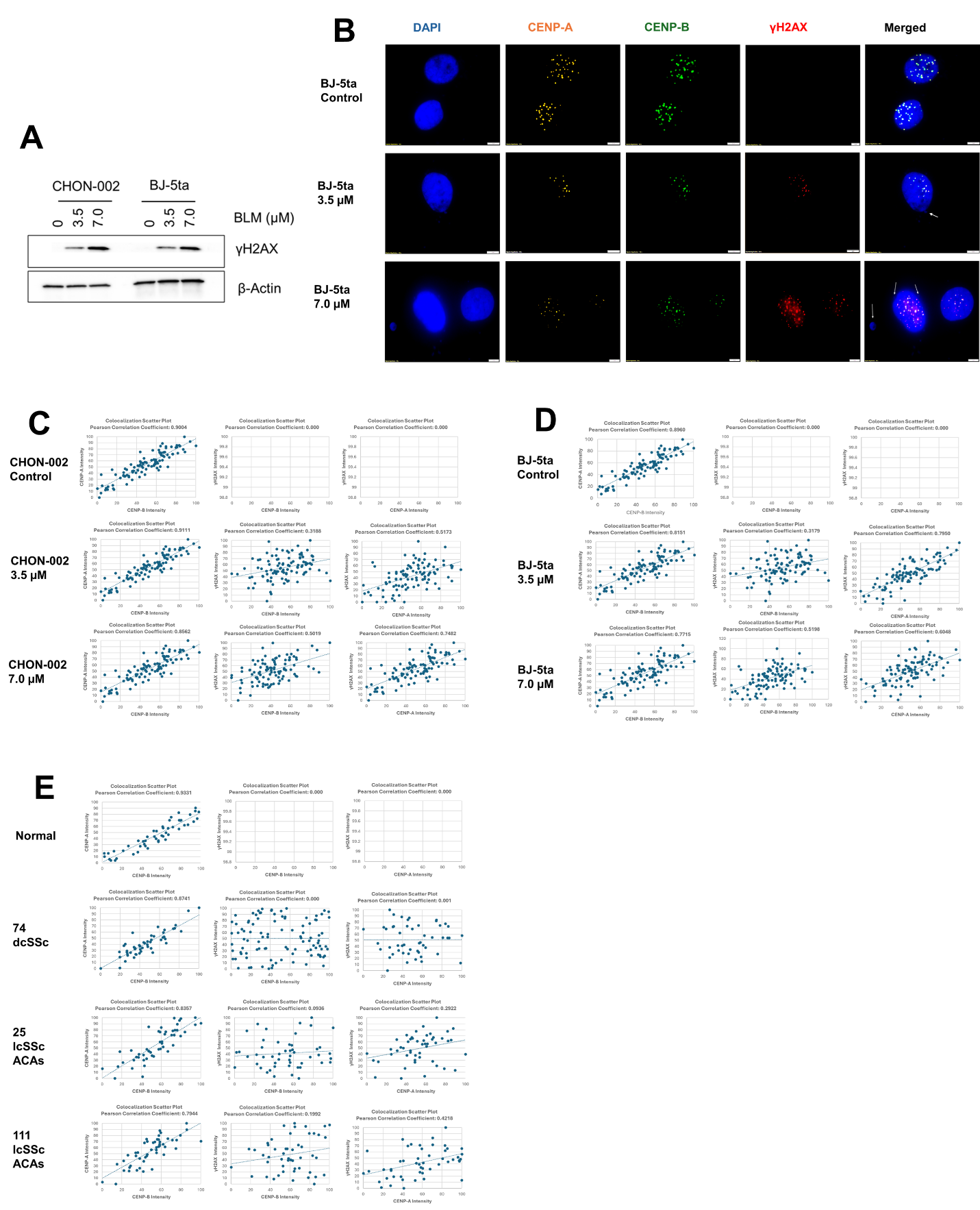

**Supplementary Fig. 3. BLM-induced DNA damage at centromere loci.** (A) Western blots showing γH2AX induction in CHON-002 and BJ-5ta fibroblasts after BLM. β-actin served as a loading control. (B) IF for BJ-5ta fibroblasts stained for CENP-A (orange), CENP-B (green), γH2AX (red), and DAPI (blue). Untreated cells showed discrete centromeric foci, whereas BLM induced strong γH2AX that localized to centromeric regions. White arrows indicate micronuclei. (C-E) Scatter plots showing colocalization between CENP-A, CENP-B, and γH2AX. Pearson’s correlation coefficients (r) were calculated from n = 50–100 cells per condition.

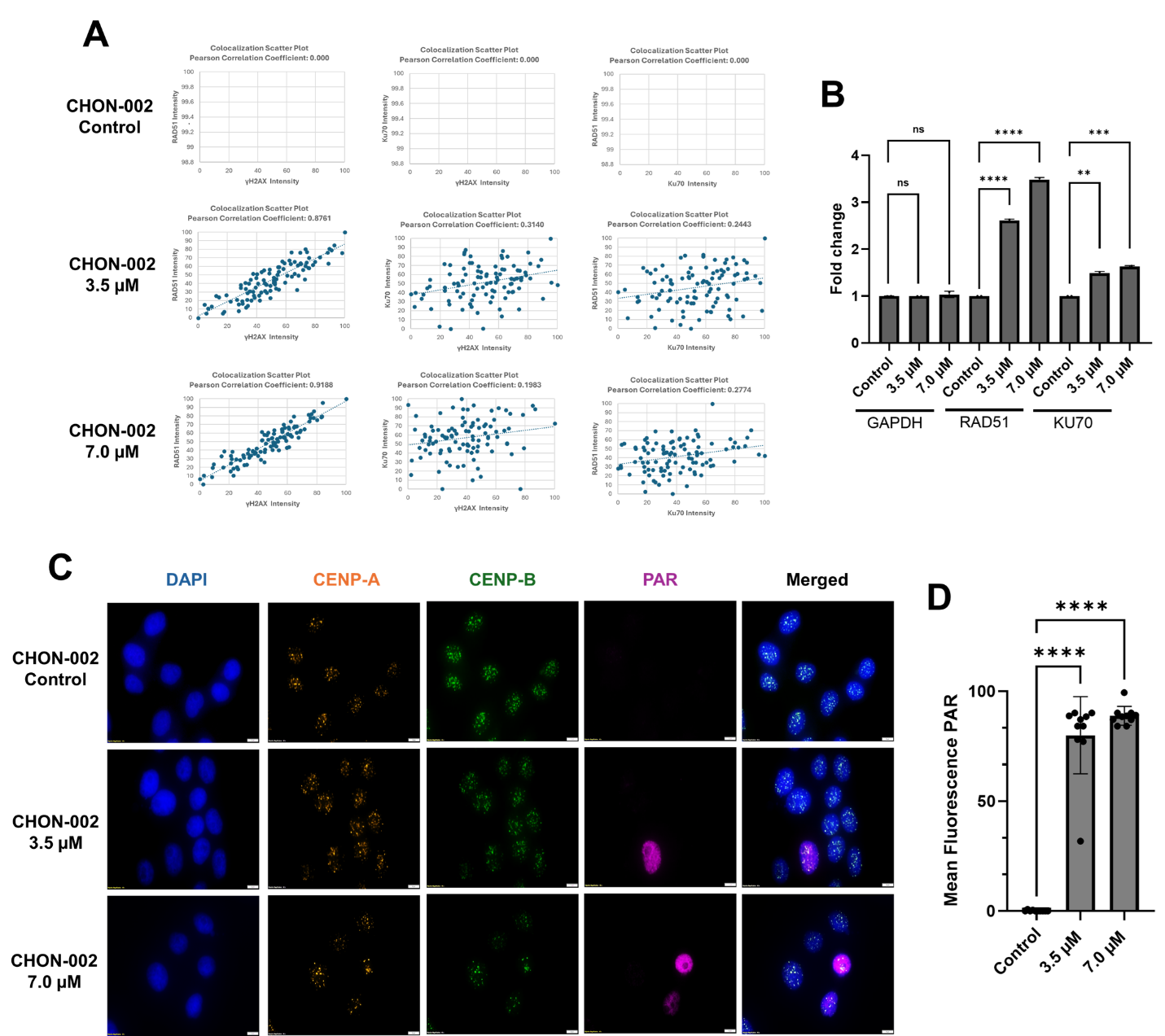

**Supplementary Fig. 4. Centromere-associated DNA repair and PAR accumulation after BLM.** (A) Scatter plots showing colocalization of RAD51, KU70, and γH2AX signals. Pearson’s correlation coefficients (r) were calculated from n = 50 cells per condition. (B) RT-qPCR analysis of RAD51 and KU70 mRNA expression after BLM versus vehicle control, normalized to GAPDH. One-way ANOVA: *****p* < 0.0001; ****p* < 0.001; ns, not significant. (C) IF for CENP-A (orange), CENP-B (green), poly(ADP-ribose) (PAR; purple) in CHON-002 fibroblasts treated with 3.5 or 7.0 µM BLM. Nuclei counterstained with DAPI (blue). Controls showed no detectable PAR, whereas BLM induced strong nuclear PAR in a subset of cells. Scale bar, 10 µm. (D) Quantification of nuclear PAR intensity in BLM-treated versus control cells (one-way ANOVA; *****p* < 0.0001, n = 10 cells per condition).

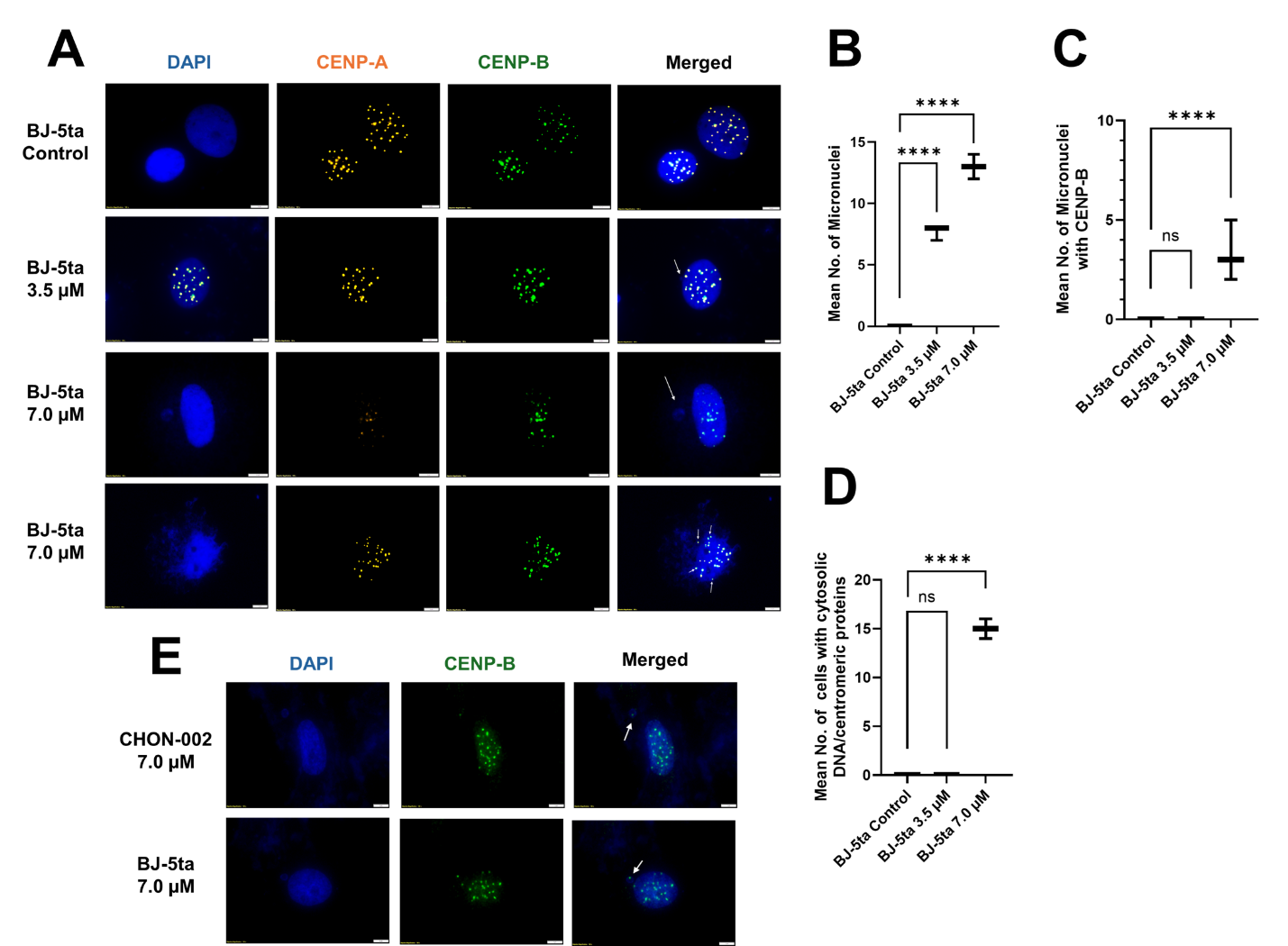

**Supplementary Fig. 5. BLM induces micronuclei and cytoplasmic centromeric foci in fibroblasts.** (A) BJ-5ta fibroblasts treated with 3.5 or 7.0 µM BLM for 3 h followed by 24 h recovery. Cells were stained for CENP-A (orange), CENP-B (green), and DAPI (blue). Untreated controls showed intact nuclei with discrete centromeric foci, whereas BLM induced micronuclei (white arrows) and cytoplasmic centromeric foci (small white arrows). Scale bar, 5 µm. (B–D) Quantification of (B) micronuclei, (C) micronuclei positive for CENP-A/B, and (D) cells with cytoplasmic centromere protein-DNA structures. Data represent mean ± SD from three independent experiments (n = 100 cells per group). One-way ANOVA, *****p* < 0.0001; ns, not significant. (E) IF for CENP-B (green) in micronuclei (white arrows) with DAPI (blue) in CHON-002 and BJ-5ta fibroblasts after 7.0 µM BLM.

| Target | Clone/Type | Host | Application/Dilution | Vendor/Cat. No. |
| --- | --- | --- | --- | --- |
| γ‑H2AX (Ser139) | 20E3 | Rabbit | IF 1: 400; WB 1:1000 | CST #9718 |
| CENP‑A | mAb | Mouse | 1:500 | MBL D115-3 |
| CENP‑B | mAb | Mouse | 1:50 | SCBT sc‑376392 |
| BANF1/BAF | EPR7668 | Rabbit | 1:100 | Abcam EPR7668 |
| MHCII DRB1 | mAb | Rabbit | 1:100 | Abclonal A7685 |
| RAD51 | mAb | Rabbit | 1:100 | Invitrogen PA5‑27195 |
| KU70 | mAb | Rabbit | 1:100 | Invitrogen PA5‑95270 |
| PAR | mAb | Mouse | 1:100 | SCBT sc‑56198 |
| Actin B | AC-15 | Mouse | 1:5000 | Sigma-Aldrich A5441 |
| Alexa Fluor-conjugated secondary antibodies (488, 555, 568, 647; Invitrogen) were used at 1:1000. | | | | |

**Supplementary Table 1. Antibodies used in this study.**
